# Computational Structural Analysis of POLG Variants R627Q and W748S with Model-Variability Controls

**DOI:** 10.64898/2026.08.25.747108

**Authors:** Andrew Friedl, Deborah Manst

## Abstract

**Background:** Comparisons between independently predicted wild-type and missense-variant protein structures can generate mechanistic hypotheses, but small apparent differences may reflect model-selection variability rather than mutation-specific effects.

**Methods:** Human mitochondrial DNA polymerase gamma (POLG; UniProt P54098) variants p.Arg627Gln (R627Q) and p.Trp748Ser (W748S) were evaluated using five AlphaFold2-PTM network-model outputs per condition generated with one random seed under matched ColabFold settings. Ten pairwise wild type comparisons at each site described between-network model-selection variability. Variant effects were summarized across five within-network wild-type-versus-variant comparisons using rotation-invariant local C-alpha pair distances and local displacement after global and local alignment. Because these comparison designs differ, the wild-type distribution was used as context rather than a mutation-effect null. Wild-type cryo-EM structure 9GGF was used for contact and interface mapping. Experimental A467T and G848S structures 9GGE and 9GGC provided contextual benchmarks.

**Results:** R627Q measurements fell within the range of between-network wild-type differences: its median mean local pair-distance change was 0.170 Å, compared with a wild-type median of 0.170 Å, and its locally aligned displacement was 0.265 versus 0.248 Å. W748S showed higher median values (0.168 versus 0.132 Å for pair-distance change; 0.236 versus 0.182 Å for locally aligned displacement), but the ranges overlapped and the comparison-design asymmetry precluded a calibrated mutation-effect percentile. Experimental A467T and G848S comparisons produced local changes of similar magnitude. In 9GGF, R627 and W748 directly shared a local microenvironment, with a minimum heavy-atom distance of 3.53 Å. R627 also formed short polar-contact candidates with D629 and D743, whereas W748 occupied a hydrophobic packing environment containing Y622 and F750. Both sites were more than 18 Å from nucleic acid, more than 30 Å from POLG2, and more than 33 Å from PZL-A in a ligand-bound structure.

**Conclusions:** Available AlphaFold2 comparisons do not establish a mutation-specific structural deformation for either variant. Experimental-structure mapping supports testable physicochemical hypotheses involving a shared R627-W748 microenvironment—loss of an arginine-centered polar network for R627Q and disruption of a buried aromatic environment for W748S—but not direct DNA, POLG2, or PZL-A contact mechanisms. Matched control substitutions and independent seeds are required to calibrate small mutation-associated structural deltas.

## Introduction

Human DNA polymerase gamma is the principal polymerase responsible for mitochondrial DNA replication and proofreading. Its catalytic POLG-encoded subunit contains an exonuclease domain, a spacer/intrinsic-processivity (IP) region, and a polymerase domain and associates with a dimeric POLG2 accessory subunit [1,2]. Disease-associated variation in POLG produces broad clinical heterogeneity, and molecular effects depend on allelic phase, biochemical context, and conformational state [3–5].

R627Q and W748S lie in the spacer/IP region rather than in the exonuclease or polymerase catalytic motifs (Figure 1). Both have been assigned to a broader functional cluster, but structural clustering does not establish a shared molecular mechanism [1]. W748S is frequently observed in cis with E1143G, and biochemical studies distinguish the isolated substitution from the W748S+E1143G protein or haplotype [3,5].

**Figure 1.**
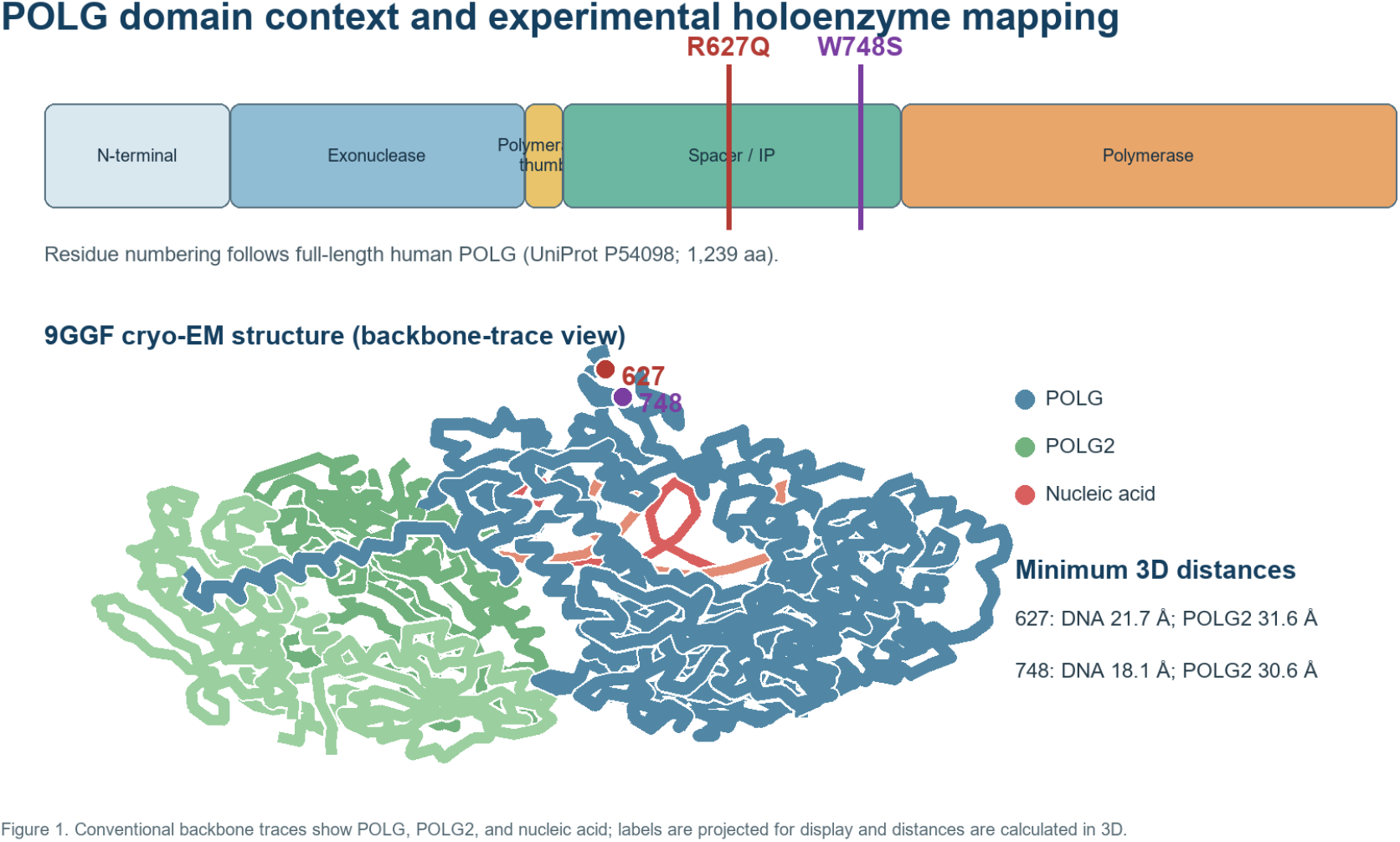
POLG domain context and experimental holoenzyme mapping.

AlphaFold2 and related methods predict protein folds with high accuracy, but the interpretation of small mutation-induced differences remains challenging [6–9]. Published studies have reported limited mutation sensitivity in some settings and useful localized strain signals in others. Consequently, a wild-type-versus-variant difference is not interpretable without a baseline reflecting variability among models generated by the same prediction process.

Therefore, this work was reframed as a calibration-aware paired analysis that asked whether R627Q or W748S structural deltas separated from available wild-type model-selection variability; whether the same descriptors were compatible with experimental A467T and G848S structures reported to preserve the general POLG architecture; and what testable local interaction hypotheses could be derived directly from an experimental wild-type holoenzyme structure.

## Methods

### Study design and claim boundary

This was a computational structural calibration study. No participant-level data, clinical outcomes, diagnostic classifications, or treatment decisions were analyzed. Structural observations were not used to classify pathogenicity or estimate mitochondrial functional reduction.

### Sequence, numbering, and prediction provenance

Residue numbering followed full-length human POLG, UniProt P54098, comprising 1,239 residues and including the mitochondrial targeting sequence. Reference, R627Q, and W748S models were generated with ColabFold version 1.6.2 (commit 0c788a0e8dca909f2c669784f9dd38e8c9682ff0) using AlphaFold2-PTM, MMseqs2 UniRef plus environmental MSAs, no templates, three recycles, one ensemble, no relaxation, no dropout, random seed 0, and five network models. Models were ranked by mean pLDDT. Complete configuration files are included in the reproducibility package.

### Model-selection variability baseline

The five wild-type network-model outputs were compared pairwise, producing ten comparisons at each variant site. This distribution measures between-network variability associated with network-model selection and ranking within a single prediction run. It is not a seed-to-seed distribution and does not capture all prediction uncertainty.

For each variant, reference and variant outputs using the same AlphaFold network-model identifier were compared, producing five within-network comparisons. Because model ranking differed across conditions, matching was based on network-model identity rather than rank number. These within-network mutation comparisons are not exchangeable with the between-network wild-type comparisons. The latter therefore provide descriptive context, not a formal mutation-effect null distribution; empirical ranks against it are not calibrated percentiles, significance tests, or error probabilities.

### Local geometric descriptors

The local neighborhood comprised residues whose wild-type C-alpha atom was within 10 Å of the variant site. The absolute change in every pairwise C-alpha distance within that neighborhood was calculated and the mean, median, and maximum were summarized. This descriptor is invariant to rotation and translation.

C-alpha displacement was evaluated after two alignments. Global alignment used all matched C-alpha atoms and can reflect domain-orientation differences. Local alignment used only neighborhood C-alpha atoms and better isolates local geometric variation. Kabsch least-squares superposition was used in both cases. Pairwise distances are descriptive and were not treated as independent statistical observations.

### Experimental benchmarks and structural context

Wild-type structure 9GGF was compared with A467T structure 9GGE and G848S structure 9GGC using the same local descriptors at residues 467 and 848. These 2.4–2.7 Å cryo-EM structures were generated in the same published study, which reported preservation of general POLG architecture [5]. They were used as contextual benchmarks rather than a formal validation set.

R627 and W748 contacts were mapped in chain A of 9GGF. Non-adjacent residues with a minimum heavy-atom distance of 4.5 Å or less were recorded. Side-chain nitrogen-to-oxygen or sulfur contacts of 3.5 Å or less were labeled polar-contact candidates; hydrogen-bond geometry, protonation, solvent, and electrostatic energy were not assigned. Minimum heavy-atom distances to nucleic acid and POLG2 were calculated in 9GGF. Distances to PZL-A (component A1IK1) were calculated in ligand-bound G848S structure 9GGB; W748 was also measured in 9GGD, whereas R627 was unresolved in that structure.

### Reproducibility and descriptive analysis

All calculations were performed with a versioned analysis script using NumPy. The output package includes the analysis code, exact configuration, source-structure identifiers, comparison-level CSV data, aggregate JSON results, figures, and a supplementary note for excluded exploratory stability predictions. Medians and ranges are reported because five variant comparisons and ten statistically dependent wild-type pairs do not support reliable parametric distribution estimates or z-scores. Cross-design empirical percentiles are not reported as calibrated mutation-effect statistics.

## Results

### R627Q values fall within between-network wild-type variation

At residue 627, ten between-network wild-type pairwise comparisons produced mean local pair-distance changes from 0.150 to 0.235 Å, with a median of 0.170 Å. Five within-network R627Q comparisons ranged from 0.111 to 0.194 Å, also with a median of 0.170 Å. Thus, the R627Q values fell within the observed between-network wild-type range.

Locally aligned displacement showed the same pattern. Wild-type comparisons ranged from 0.184 to 0.384 Å with a median of 0.248 Å; R627Q comparisons ranged from 0.206 to 0.348 Å with a median of 0.265 Å. The distributions overlapped completely. Global-alignment displacement was also centered within wild-type variability, illustrating that a single-model displacement can overstate evidence for a mutation-specific effect.

### W748S medians exceed corresponding wild-type medians, with overlapping ranges

At residue 748, between-network wild-type pair-distance changes ranged from 0.106 to 0.207 Å with a median of 0.132 Å. Within-network W748S comparisons ranged from 0.161 to 0.237 Å with a median of 0.168 Å. The W748S median was higher, but the ranges overlapped.

Locally aligned displacement ranged from 0.157 to 0.315 Å among between-network wild-type comparisons and from 0.203 to 0.367 Å for within-network W748S comparisons, with medians of 0.182 and 0.236 Å (Figure 2). The distributions overlapped. In contrast, displacement after global alignment was lower for W748S than for the wild-type comparison median, confirming that global and local alignment measure different sources of variation and should not be used interchangeably.

**Figure 2.**
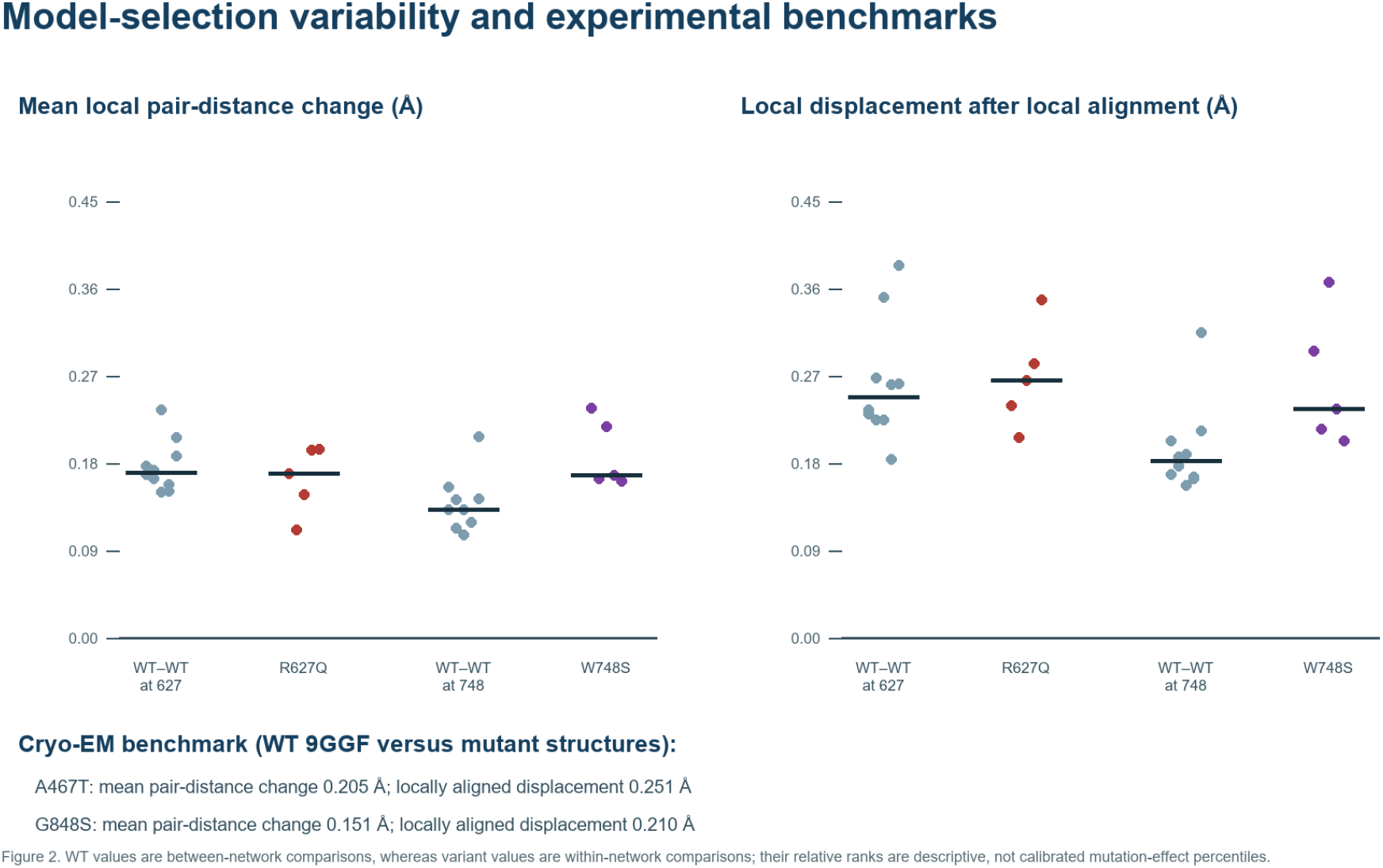
Model-selection variability and experimental benchmarks.

Site predicted Local Distance Difference Test (pLDDT), AlphaFold’s per-residue structural confidence score, also varied across network models. At R627, wild type pLDDT ranged from 72.25 to 83.06 and R627Q from 72.00 to 82.81. At W748, wild type values ranged from 80.44 to 89.00 and W748S from 74.88 to 89.69. The direction of the variant-minus-reference pLDDT difference was not consistent across W748S network models.

### Experimental comparisons have similar local-delta magnitudes

Comparison of experimental wild-type 9GGF with A467T 9GGE produced a mean local pair-distance change of 0.205 Å and a locally aligned displacement of 0.251 Å. The G848S 9GGC comparison produced values of 0.152 and 0.210 Å. These values are similar in magnitude to both the AlphaFold wild-type variability and variant comparisons. This is consistent with the published conclusion that A467T and G848S preserve general architecture while allowing localized differences [5]. The wild-type and mutant atomic models were all initialized by docking the same 4ZTZ structure into their respective cryo-EM maps before real-space fitting [5]. Shared starting-model bias could reduce apparent differences, whereas independent reconstruction and refinement variation could increase them. Accordingly, the benchmark supports the narrower conclusion that sub-angstrom descriptor values should not be interpreted as evidence of functional disruption by magnitude alone.

### Experimental 9GGF supports specific local hypotheses

R627 and W748 are themselves nearest-neighbor members of the same local microenvironment: their minimum heavy-atom distance in 9GGF was 3.53 Å. This proximity links the two site-centered contact maps but does not by itself demonstrate energetic coupling. A prespecified R627Q/W748S double-mutant experiment could test whether their biochemical effects are additive or interacting. R627 formed short side-chain polar-contact candidates with D743 (2.70 Å) and D629 (2.71–3.15 Å) in 9GGF. Replacement of arginine with glutamine removes a positive charge and could alter this local interaction network (Figure 3). The calculation identifies a falsifiable contact hypothesis but does not establish the persistence, energetic contribution, or functional importance of these interactions.

**Figure 3.**
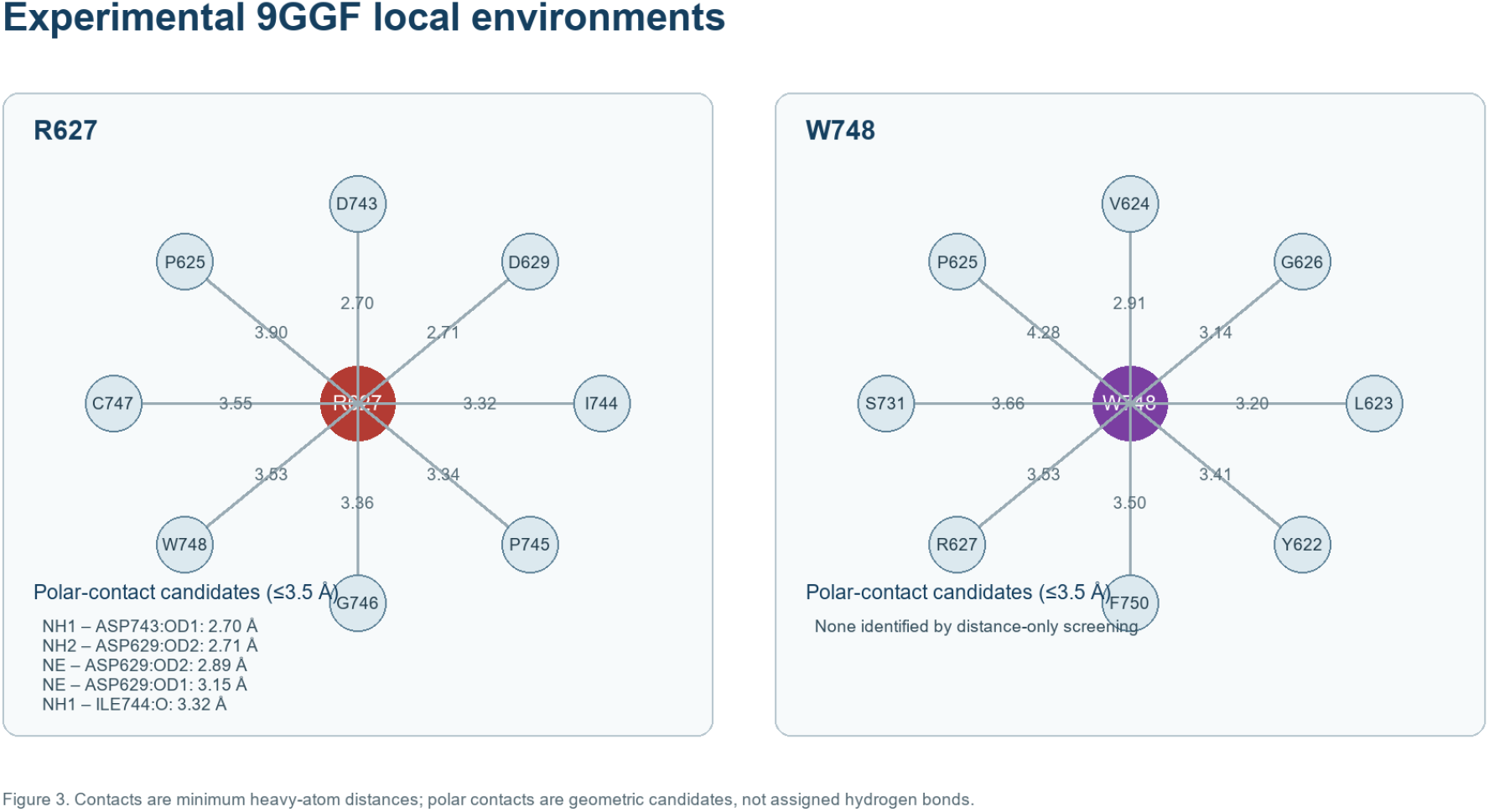
Experimental local environments of R627 and W748.

W748 occupied a nonpolar and aromatic environment. Its nearest non-adjacent contacts included V624 (2.91 Å), G626 (3.14 Å), L623 (3.20 Å), Y622 (3.41 Å), F750 (3.50 Å), and R627 (3.53 Å). No side-chain polar-contact candidate was identified for W748 by the distance-only rule. Replacement of tryptophan with serine therefore creates a plausible loss-of-aromatic-packing hypothesis independently of the non-separating AlphaFold deformation measurements.

### Neither site forms a direct DNA, POLG2, or PZL-A contact

In 9GGF, R627 was 21.70 Å from the nearest nucleic-acid atom and 31.62 Å from POLG2. W748 was 18.14 Å from nucleic acid and 30.65 Å from POLG2. In PZL-A-bound 9GGB, R627 and W748 were 37.61 and 33.69 Å from the ligand, respectively; the W748 distance in 9GGD was 34.10 Å. These measurements do not support direct-contact mechanisms involving DNA, POLG2, or the PZL-A pocket.

### Summary of contextual descriptors

| Descriptor | R627Q | W748S |
| --- | --- | --- |
| Variant mean pair-distance change, median (range), Å | 0.170 (0.111–0.194) | 0.168 (0.161–0.237) |
| Site-specific WT pair-distance baseline, median (range), Å | 0.170 (0.150–0.235) | 0.132 (0.106–0.207) |
| Variant locally aligned displacement, median (range), Å | 0.265 (0.206–0.348) | 0.236 (0.203–0.367) |
| Site-specific WT displacement baseline, median (range), Å | 0.248 (0.184–0.384) | 0.182 (0.157–0.315) |
| Nearest nucleic acid in 9GGF, Å | 21.70 | 18.14 |
| Nearest POLG2 in 9GGF, Å | 31.62 | 30.65 |
| Nearest PZL-A in 9GGB, Å | 37.61 | 33.69 |

## Discussion

The contextual analysis changes the interpretation of both variants. R627Q pair-distance and displacement measurements fell within the observed between-network wild-type range. W748S produced higher median local metrics, but its ranges overlapped the corresponding wild-type ranges. These observations support targeted follow-up rather than a claim of mutation-specific structural deformation.

The contrast between global and local alignment explains why individual displacement estimates can differ across analytical approaches. Global displacement was strongly affected by variation outside the mutation neighborhood, whereas locally aligned displacement and pair-distance changes focused on local geometry. The metrics therefore need not track each other. For mutation interpretation, the locally aligned and rotation-invariant descriptors are more directly relevant, but they still require an empirical variability baseline.

The experimental A467T and G848S comparisons add an important contextual point. Both produced local descriptor magnitudes comparable to the AlphaFold comparisons even though the published structural analysis reported no change in general architecture. Shared initialization from 4ZTZ could reduce apparent differences, while independent reconstruction and refinement variation could increase them. G848S nevertheless creates a specific local hydrogen bond to the V845 backbone carbonyl [5]. Thus, small local differences can coexist with preserved architecture, and useful mechanistic interpretation depends on named interactions and experimental context rather than a displacement threshold alone.

Experimental 9GGF mapping supports more specific hypotheses for R627Q and W748S. The 3.53 Å minimum heavy-atom distance places R627 and W748 in a shared local microenvironment. R627 is positioned in an arginine-centered polar network involving D629 and D743, which R627Q could weaken or reorganize. W748 is surrounded by hydrophobic and aromatic contacts, including Y622 and F750, making loss of aromatic packing plausible after substitution by serine. The proximity motivates a double-mutant interaction experiment, but it does not establish coupling or non-additivity. These hypotheses arise from the wild-type experimental environment and residue chemistry; they are not proven consequences of the predicted variant models.

The 2025 biochemical study provides complementary functional context. All four tested mutant proteins showed reduced processivity, while W748S was the exception that did not show reduced template affinity in the stalled elongating condition [5]. Importantly, the recombinant W748S protein contained E1143G in cis. Those findings support an indirect and context-dependent defect but cannot be attributed to isolated W748S alone.

### Limitations

The principal limitation is the asymmetry between the descriptive wild-type context and the variant comparison. The wild-type range contains between-network comparisons, whereas each variant delta is a within-network wild-type-versus-variant comparison. These quantities are not exchangeable, so their overlap or relative rank cannot establish a mutation-effect null result. The model set also contains five AlphaFold network models from one seed, not independent prediction seeds, and pairwise comparisons among shared models are statistically dependent.

The A467T and G848S structures differ from wild type by mutation and by experimental reconstruction, so their differences are contextual benchmarks rather than pure mutation effects. Contact screening did not assign bond angles, protonation, solvation, dynamics, or interaction energies. The analysis includes only four variant sites, no matched benign control-substitution panel, and no biochemical measurements generated in this study.

### Next validation steps

A prespecified extension should first generate a within-network control-substitution null using conservative surface substitutions or independently justified population-observed POLG variants processed with the identical pipeline. It should also generate at least ten independent seeds for wild type, controls, and each study variant while holding MSA, templates, recycles, and software version constant. Control selection and analysis should be blinded to structural outcomes where feasible. Conclusions should be based on matched-comparison calibration, out-of-seed replication, and interaction-level agreement with experimental structures rather than on a single displacement threshold.

## Conclusions

R627Q values fell within the observed between-network wild-type range, while W748S produced higher medians with overlapping ranges. Experimental structures identify testable local mechanisms in a shared R627-W748 microenvironment: disruption of an R627-centered polar network and loss of the W748 aromatic packing environment. Neither variant site directly contacts DNA, POLG2, or the PZL-A pocket in the analyzed structures. These findings support focused biochemical and double-mutant experiments while demonstrating that small wild-type-versus-variant structure differences are insufficient for mechanistic or clinical classification without matched controls.

## Data and Code Availability

The reproducibility package is permanently archived on Zenodo under a CC BY 4.0 license [10]. It contains the analysis script, exact ColabFold configuration, model and score files, comparison-level data, aggregate results, figures, source-structure manifest, and a supplementary note containing the excluded exploratory DynaMut2 values. No participant-level data were used or generated.

## Ethics Statement

This computational study used protein models, publicly available experimental structures, and published literature. It did not involve human participants, identifiable participant-level data, animals, or biological specimens; institutional review board approval and informed consent were therefore not required.

## Author Contributions

Andrew Friedl conceived the study, developed the analysis workflow, interpreted the results, and drafted the manuscript with AI-assisted editorial support. Deborah Manst contributed to interpretation and critical revision of the manuscript.

## AI Assistance Disclosure

OpenAI’s ChatGPT/Codex was used as an AI-assisted coding, analysis, writing, organization, and editorial tool. Anthropic’s Claude was used as an AI-assisted manuscript reviewer to critique the methods, interpretation, and presentation. These AI systems are not authors or formal peer reviewers. The human authors remain responsible for the accuracy, originality, interpretation, and integrity of the manuscript.

